# Thalamic-medial temporal lobe connectivity underpins familiarity memory

**DOI:** 10.1101/665687

**Authors:** Alex Kafkas, Andrew R. Mayes, Daniela Montaldi

## Abstract

The neural basis of memory is highly distributed, but the thalamus is known to play a particularly critical role. However, exactly how the different thalamic nuclei contribute to different kinds of memory is unclear. Moreover, whether thalamic connectivity with the medial temporal lobe (MTL), arguably the most fundamental memory structure, is critical for memory, remains unknown. We explore these questions using an fMRI recognition memory paradigm that taps familiarity and recollection (i.e., the two types of memory that support recognition) for objects, faces and scenes. We show that the mediodorsal thalamus (MDt) plays a material-general role in familiarity, while the anterior thalamus plays a material-general role in recollection. Material-specific regions were found for scene familiarity (ventral posteromedial and pulvinar thalamic nuclei) and face familiarity (left ventrolateral thalamus). Critically, increased functional connectivity between the MDt and the parahippocampal (PHC) and perirhinal cortices (PRC) of the MTL underpinned increases in reported familiarity confidence. These findings suggest that familiarity signals are generated through the dynamic interaction of functionally connected MTL-thalamic structures.

## Introduction

The ability to discriminate previously encountered stimuli from new ones is critical to our efficient processing of information. A sense of familiarity allows the identification of information as old, potentially triggering further memory search, while the detection of novelty prompts the encoding of new information into memory. Familiarity memory is a type of recognition memory that is derived from the comparison between incoming information and previously stored representations, but it remains to be established how the brain generates familiarity signals as a function of stored representations and whether these signals are integrated across different brain regions. Indeed, across many functional neuroimaging studies (e.g., Montaldi et al. 2006; Skinner and Fernandes 2007; Kafkas and Montaldi 2012, 2014; Pergola and Suchan 2013; Scalici et al. 2017), familiarity memory has been shown to engage a number of brain regions, which suggests that several interacting processes support familiarity memory and that these processes engage functionally connected brain regions. An important endeavor, therefore, is to characterize the role of the dynamic interactions of networks involved in memory and the integration of familiarity signals.

The prominent role of the MTL regions, including the hippocampus, the perirhinal (PRC), entorhinal (ERC) and parahippocampal (PHC) cortices, in supporting recognition memory is clearly evident from many neuroimaging and neuropsychological studies (Aggleton and Brown 1999; Eichenbaum et al. 2007; Montaldi and Mayes 2010; Kafkas and Montaldi 2012; Kafkas et al. 2017). However, the structures of the MTL do not function in isolation, and recent studies provide further evidence for the role of the thalamus and its specific nuclei in supporting recognition memory (for reviews see Aggleton et al. 2011; Mitchell and Chakraborty 2013; Carlesimo et al. 2015). In the influential model proposed by Aggleton and Brown (1999; Aggleton et al. 2011), two functional systems were identified: one which supports recollection and recall-like memory, and is often referred to as the “extended hippocampal system” as it includes the hippocampus and the anterior thalamus (ANt), as well as the fornix, mammillary bodies and mammillothalamic tract, which connect them. The other, which supports familiarity memory, includes the PRC and the mediodorsal thalamus (MDt), which are connected through the amygdalofugal pathway. Although this model ascribes a prominent role to at least two major thalamic regions (ANt and MDt), their specific roles within the respective functional systems are poorly understood. Neuropsychological evidence supports the idea that their role cannot be merely assistive as dense anterograde amnesia can follow damage to the ANt and the MDt (Kishiyama et al. 2005). However, whether damage in only one of these two structures (and especially the MDt) is sufficient to generate anterograde amnesia is still debated (Mitchell and Chakraborty 2013; Carlesimo et al. 2015) and perhaps it depends on the kind of memory tasks used. For example, if MDt has a selective role in familiarity then damage in this structure will leave recall/recollection unaffected and therefore the observed anterograde amnesia will be much milder than if damage also includes the ANt.

Despite evidence supporting a degree of functional specialization within the thalamus, neuropsychological and neuroimaging studies have produced conflicting findings (Kafkas and Montaldi 2012, 2014; Pergola et al. 2012, 2013; Edelstyn et al. 2016; Danet et al. 2017). In particular, the majority of neuropsychological evidence shows that the MDt supports familiarity memory (Edelstyn et al. 2016; Newsome et al. 2018); nevertheless, some evidence that it supports both familiarity and recollection also exists (Danet et al. 2017). The apparent similarity between the MTL and MDt patterns of specialization suggests that strong interplay is likely, however, to date, there is no direct evidence of a functional interplay, or coupling, between MDt and the MTL cortices when participants make recognition (or familiarity) decisions. The aim of the present study, therefore, is to further characterize the contribution of the thalamic nuclei to familiarity and/or recollection, and to determine the extent of their functional coupling with the structures of the MTL, while people engage in recognition memory.

In a recent fMRI study (Kafkas et al. 2017) we focused on the functional heterogeneity of the MTL and using three types of pictorial stimulus (objects, faces and scenes), measured the MTL activation patterns that accompanied familiarity and recollection responses as a function of stimulus type. The study showed that familiarity-driven activations within the MTL cortical regions were sensitive to the type of stimulus in memory, with PRC and ERC responding to object familiarity, PHC responding to object and scene familiarity and the amygdala responding to face familiarity. In contrast, the hippocampus was not found to respond to familiarity but to recollection, and critically in a material-general fashion.

Here we present a further analysis of the data from Kafkas et al. focusing on examining the roles played by the different thalamic nuclei in supporting familiarity and/or recollection memory, and asking whether these roles are material-specific (i.e., responding selectively to one stimulus category) or material-general (i.e., responding across the three types of pictorial stimuli). Specifically, based on previous theoretical models and findings (Aggleton and Brown 1999; Kafkas and Montaldi 2012, 2014), we hypothesised that the MDt is sensitive to familiarity. However, the degree to which this activation applies to all types of stimuli remains unknown and is investigated in the present study. Furthermore, connectivity analysis is used to explore the extent to which thalamic regions sensitive to familiarity or recollection, are functionally coupled with MTL structures during a recognition memory task. We hypothesised that connectivity between the MDt and the neocortical areas of the MTL and the amygdala increases when familiarity is reported. In contrast, we hypothesised that the anterior thalamus will be functionally connected with the hippocampus when participants report instances of recollection (relative to strong familiarity). This investigation is central to understanding the role the thalamic nuclei play within the wider memory system and, critically, for the first time, explores the degree to which thalamic regions (and especially the MDt) work together with MTL structures to support recognition memory.

## Materials and Methods

### Participants

Data from 17 right-handed participants (5 females) with a mean age of 23.30 years (SD = 3.40) were used in the analyses reported here. From an original sample of 20 participants, three participants were excluded from further analysis (two participants due to excessive movement in the MRI scanner and one due to chance memory performance in the recognition task). Healthy volunteers without any self-reported neurological or psychiatric disorder were recruited from the student population of the University of Manchester and the general public. All had normal or corrected-to-normal vision and all received £20 after their participation in the study. The procedures adopted in the study and the consent procedure had been approved by the National Research Ethics Service (North West-GM South) before the study commenced.

### Materials and Procedure

The experiment used 420 colour stimuli (15 for practice: five per stimulus type), consisting of 140 objects (man-made and natural), 140 faces (male and female) and 140 scenes (landscape images). Before scanning, participants encoded a series of 90 objects, 90 faces and 90 scenes presented in 10 alternating blocks of 9 stimuli each (i.e., 270 stimuli at encoding in total). A shallow encoding procedure was adopted in which participants were asked to make matching-to-sample decisions for each stimulus. In this task, each object, face and scene trial was presented as a triplet of 3 very similar pictures of the same stimulus and participants were asked to indicate which of the lower two pictures matched the target presented on top. In each case one of the two lower pictures had been minimally modified in size relative to the target (for objects and faces). In the case of the scenes, the modified picture had been slightly shifted horizontally (either leftwards or rightwards) by a few millimeters. The allocation of the modified foil in each trial (either lower left or lower right) was randomly determined and participants were given 4s to select the matching picture, which they did by pressing “1” for left or “0” for right using a PC keyboard. This encoding task has been extensively piloted and has been used in previous papers as it generates a predominance of familiarity-based recognition later at retrieval (Kafkas and Montaldi 2012, 2014, 2015; Kafkas et al. 2017).

The retrieval phase was completed inside the MRI scanner, approximately 25-30 minutes after the encoding phase. This gap included preparation of each participant to enter the MRI scanner, instructions on how to make the memory decisions (see Suppl. Material), completion of two practice blocks and acquisition of the T1 image. In the retrieval phase participants were asked to select whether each presented stimulus was familiar (F), recollected (R) or new (N). Familiarity decisions were made using a 3-point scale of increasing strength (F1 = weak, F2 = moderate and F3 = strong familiarity), while two additional options were provided for R and N responses. For each stimulus, participants were asked to indicate the strength of their feeling of familiarity if they thought that the item was familiar, or select R or N. Participants were instructed to use R only when they had spontaneously recollected non-stimulus associative information regarding the previous presentation of the stimulus at encoding. Critically, participants were not asked to effortfully recollect but to report any spontaneous recollections. An R response therefore, could be a thought or any other extraneous information associated with the stimulus at encoding (e.g., they recall, from the study episode, that they had likened the bicycle stimulus to their own bicycle, or they recall that the stimulus was presented first/last or that they sneezed when the stimulus was on the screen). The distinction between F and R (see Kafkas and Montaldi 2012; Migo et al. 2012) was explained in detail before commencing the retrieval block in the MRI scanner and participants were asked to provide examples from their own experience of memory to illustrate that they had understood the distinction (for the F/R instructions see Suppl. Material). Also, the procedure was practiced twice, once outside the scanner and once inside while the T1 anatomical image was acquired. In the first practice block participants were asked to justify any R responses to check adherence with the F/R instructions and appropriate corrective feedback was provided.

At retrieval 405 stimuli were presented consisting of 270 studied stimuli and 135 new foils. This means that for each type of pictorial stimulus participants were presented with 90 studied stimuli plus another 45 new foils. A pseudo-random sequence was adopted for presenting objects, faces and scenes at retrieval. Blocks of 15 stimuli were presented in an alternating fashion with each block containing one type of stimulus. Null events (baseline fixation trials) were also presented within each block (3 null events in each block; i.e., a total of 81 null events). Each stimulus appeared for 3s followed by a 1s fixation cross and participants indicated their decision by selecting 1 of the 5 available buttons on an MR-compatible response box. Three buttons on one hand were allocated to the F responses, while 2 buttons on the other hand were used for the other two responses (R and N). The allocation of these buttons to left and right hand was counterbalanced across participants.

### fMRI acquisition and pre-processing

Data were acquired on a 3T Philips (Achieva) scanner using a gradient echo-planar sequence. For the functional data the blood oxygenation level dependent (BOLD) contrast was used and 840 volumes (40 slices each) were collected for each participant covering the whole brain (TR = 2.5s; TE = 35ms; voxel size = 2.5 × 2.5 × 3.5 mm). Anatomical images (T1) were also collected, for each participant, at the beginning of the scanning session with a matrix size 256 × 256 giving 180 slices and voxel size of 1mm isotropic.

Data pre-processing and analyses were conducted using SPM8 (Statistical Parametric Mapping, Wellcome Trust Centre for Neuroimaging, http://www.fil.ion.ucl.ac.uk/spm/) and data quality was examined using the ArtRepair software (http://cibsr.stanford.edu/tools/human-brainproject/artrepair-software.html). Linear interpolation to the adjacent slices was implemented in ArtRepair to correct major artefacts in the individual time-series and this was used for less than 5% of the slices from two subjects. The time-series of the other 15 subjects in the sample did not suffer from any major artefacts and were not further corrected. The pre-processing steps in SPM included registration of the functional time-series to the mean image (using six-parameter rigid body transformation), reslicing using sinc interpolation and slice-time correction (to the middle slice) to account for slice acquisition time differences in the functional volumes. The anatomical scans of each individual were coregistered to the mean EPI image for each participant and spatial normalization of the EPI and T1 images was performed using DARTEL as implemented in SPM8 (Ashburner 2007). The functional data were resliced to 3mm isotropic volumes and were minimally spatially smoothed with a 4mm full width half maximum (FWHM) Gaussian kernel.

### First-level and second-level whole brain analyses

After preprocessing, a general linear model (GLM)-based analysis was implemented to analyze the EPI data at the first (subject) level. As standard in event-related functional data, a canonical hemodynamic response function (Friston et al. 1998) was convolved with a series of delta functions corresponding to the onset of each event (i.e., the onset of each trial). For each individual, two models were specified, a parametric and a categorical model, each including all response outcomes for the three types of stimulus used in the experiment. Additional regressors of no-interest included trials with no behavioural response, the six movement parameters produced at realignment and residual movement artefacts from ArtRepair. The inclusion of response times (RTs) as regressors (to account for variance explained by the variability in RTs characterizing each trial) did not produce any change in the main findings, consistent with our previous studies with similar methodology (e.g., Kafkas and Montaldi 2012, 2014; Kafkas et al. 2017; Mayes et al. 2019). Finally, the data were high-pass filtered using a cut-off of 128s.

Parametric analysis was performed to isolate the brain regions that show sensitivity in their activity for increases or decreases in reported familiarity strength for scenes, objects and faces. The performed analysis steps are in accordance with the standard parametric analysis methodology in SPM (Büchel et al. 1998). Familiarity hits were included in the parametric model for each stimulus type and the reported familiarity strength (F1, F2, F3) was convolved, as a covariate, with the stimulus specific haemodynamic response function (HRF). An additional level of zero familiarity (F0) was also included in the parametric model for misses (i.e., old items that were reported as new). At the first-level, parametric *t*-contrasts were created separately for each stimulus type and were then used in the group (second-level) analysis. The parametric contrasts modeled linear (or monotonic) increases and decreases in activity as a function of reported familiarity. Non-linear parametric effects were also included in the model but as these produced no extra activation (to those identified in the linear contrasts), are not reported separately.

As the aim of the analyses was to isolate both material-general familiarity brain responses and material-specific effects, a conjunction methodology was combined with the application of exclusive masks. Therefore, shared neural familiarity effects were produced by means of a conjunction analysis across the three parametric contrasts for scenes, objects and faces (Friston et al. 2005). This conjunction method isolated the brain regions that are consistently activated as a function of reported familiarity for all three types of stimuli. Both parametric activations (F0 → F3) and parametric deactivations (F3 → F0) across familiarity strength were examined, but only areas responding to increases in familiarity strength were found (see Results). Material-specific familiarity responses were explored by using the separate parametric contrasts for each stimulus type exclusively masked (at *P* < 0.05) by the parametric effects of the other two types of stimuli. Therefore, whole-brain scene-specific familiarity effects are reported when they survived exclusive masking by the parametric responses to objects and faces. Similarly, object-specific familiarity effects survived exclusive masking by face and scene familiarity, while face-specific familiarity effects survived exclusive masking by object and scene familiarity activations. Finally, R responses were contrasted with F3 responses (R > F3 contrast) to reveal brain regions that respond to recollection. The “Thalamus Atlas” (Krauth et al. 2010) was used to classify the location of the activations in the different thalamic nuclei.

### PPI connectivity analyses

Psychophysiological interaction (PPI) analyses were also performed in SPM8 (Friston et al. 1997). PPI analyses were used to explore the connectivity patterns between the identified thalamic regions and the structures of the MTL or the whole brain, as a function of reported familiarity. According to this method, the modulatory effect of the activity of one brain region on another is explored under the influence of a psychological contrast, in this case the strength of reported familiarity. The two regions, the seed area and the target (identified) region, are thought to be functionally coupled when specific experimental contrasts are examined. Therefore, a significant PPI in the present study indicates that activity in a target region is modulated by or co-varies with activity in the seed area when participants experience familiarity memory for the three types of stimuli. As this study was interested in examining the functional coupling of the thalamus with other important regions of the recognition memory network, four thalamic regions were selected as seed areas and their functional coupling was examined in four PPI analyses. In the main parametric/whole-brain analysis, three thalamic regions were found to respond to stimulus familiarity, either collectively across the three types of stimuli (mediodorsal thalamus, MDt; MNI: −6 −9 6), specifically for face (ventrolateral thalamus, VLt; MNI: −12 −15 0) or specifically for scene familiarity (ventral posteromedial thalamus, VPt; MNI: 9 −21 −3). Finally, an anterior thalamic region (ANt; MNI: 6 −8 12), that was found to respond to recollection (R) more than strong familiarity (F3), was also used as a seed area. The size and the location of these seeds are shown in Suppl. Figure 2.

To perform the PPI analyses a new design matrix was created for each individual and each seed region. For each participant, the deconvolved functional time-series from a 6mm radius sphere, within the group local maxima, were extracted from each seed thalamic region. The PPI GLM matrix for each seed included a) the time-series from the seed region (physiological variable), b) the psychological contrast of interest (psychological variable) and c) an interaction term expressed as the product between the physiological and psychological variables. After estimating the GLM with these regressors, *t*-contrasts of the interaction term were created at the first-level analyses and were then analyzed at the group level using one-sample *t*-tests (Friston et al. 1997). Both positive and negative PPIs were examined, as the parametric psychological contrasts used in the PPI analyses (see previous section about the parametric analyses) allowed the exploration of increased coupling between seed and targets for increases in familiarity strength (positive PPI), but also increased coupling between seed and targets for decreases in reported familiarity strength (negative PPI). Significant task-related negative PPIs were only isolated in the case of face familiarity (VLt seed). Conventionally, 6mm spheres (or bigger) have been used in PPI analyses to extract time-series from seed regions, but due to the small size of the regions of interest (thalamic seeds) the PPI analysis steps were also repeated for 3mm spheres. These produced qualitatively similar results and can be seen in Suppl. Table 9.

### Thresholding, correction for multiple comparisons and behavioural analyses

Significant clusters of activation in the whole-brain univariate and the PPI analyses were determined using nonparametric permutation testing (with 5000 permutations) for the targeted contrasts implemented in the Statistical NonParametric Mapping toolbox (SnPM 13; URL: http://warwick.ac.uk/snpm). Using this method, activations are reported as significant if they survived a family wise error (FWE) corrected level of *p* < 0.05 (at an initial voxel-wise *p* < 0.001). This procedure was informed by recent evidence that commonly used parametric cluster-based correction methods result in inflated type I error (Eklund et al. 2016), whereas nonparametric permutations have been proposed to provide a conservative estimate of significant effects with very few assumptions (Nichols and Holmes 2002; Eklund et al. 2016). For the analysis of the behavioural data (accuracy and RTs) non-parametric tests were used (Friedman’s ANOVA for repeated measures and Wilcoxon signed-rank test for post-hoc contrasts with Bonferroni correction). It should be noted that the effects were exactly the same when parametric tests were employed.

## Results

The behavioural results are illustrated in Figure 1b and the proportion of trials across all response outcomes are reported in Suppl. Table 1. The behavioral analyses revealed differential levels of accuracy across familiarity strength (*χ*^2^(2) = 33.09, *p* < 0.001) and across stimulus type (*χ*^2^(2) = 19.01, *p* < 0.001). As expected, post-hoc analysis with Wilcoxon signed-rank test revealed greater accuracy for increased levels of familiarity [F3 (M = 0.88, SD = 0.09) > F2 (M = 0.71, SD = 0.13) > F1 (M = 0.50, SD = 0.11), all *p*s < 0.001], while objects (M = 0.72, SD = 0.14) were characterized by greater familiarity accuracy than faces (M = 0.66, SD = 0.09; *p* = 0.001). No other difference in familiarity accuracy across the three types of stimuli was significant. Finally, recollection accuracy (M = 0.90, SD = 0.13) was closely matched to F3 accuracy (M = 0.90, SD = 0.09; *Z* = 0.85, *p* = 0.40). A similar analysis of RTs revealed, significant differences across familiarity strength (*χ*^2^(2) = 10.71, *p* = 0.005) with decreased RTs characterizing more confident responses (F3 faster than F2 and F1, *p*s < 0.003). Also the significant main effect of stimulus type (*χ*^2^(2) = 25.76, *p* < 0.001) indicated longer RTs for scenes (M = 1943 ms, SD = 42.61 ms) than faces (1656 ms; SD = 56.43 ms) and objects (1620 ms; SD = 39.52 ms; both *p*s < 0.001). Matched RTs characterized F3 and R responses across all stimulus types collapsed: (*Z* = 0.74, *p* = 0.46) and separately for each type of stimulus (Scenes F3 vs R: *Z* = 1.01, *p* = 0.31; Objects F3 vs R: *Z* = 0.71, *p* = 0.48).

**Figure 1.**
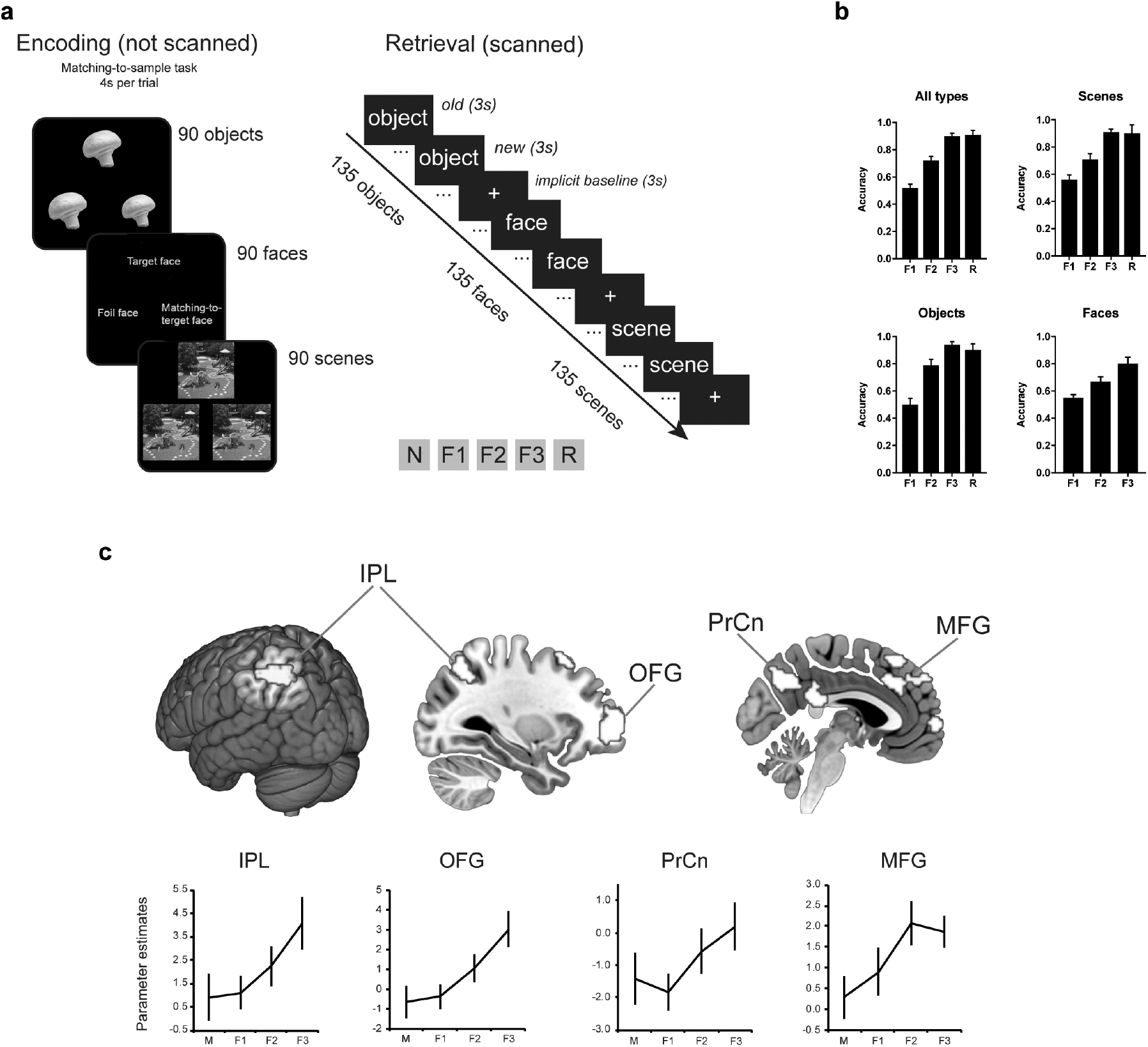
Experimental design and whole-brain material-general familiarity findings. a) Experimental design of the fMRI study. Participants (n = 17) encoded series of objects, faces and scenes using a shallow (matching-to-sample) encoding task. At retrieval in the MRI scanner they were asked to rate feelings of familiarity (F1 = weak; F2 = moderate; F3 = strong familiarity), to detect new stimuli (N) and report instances of spontaneous recollections (R). b) Accuracy across familiarity and recollection responses collapsed for all stimulus types and separately for scenes, objects and faces. Faces did not generate enough R responses and therefore only F responses are reported. c) Whole-brain material-general familiarity activations and parameter estimate plots. Abbreviations: M = misses; IPL = Inferior parietal lobe; OFG = Orbitofrontal gyrus; PrCn = Precuneus; MFG = Middle frontal gyrus. Error bars show the standard error of the mean. Note: Face stimuli are not shown in 1a in accordance with bioRxiv policies regarding pictures of faces.

**Figure 2.**
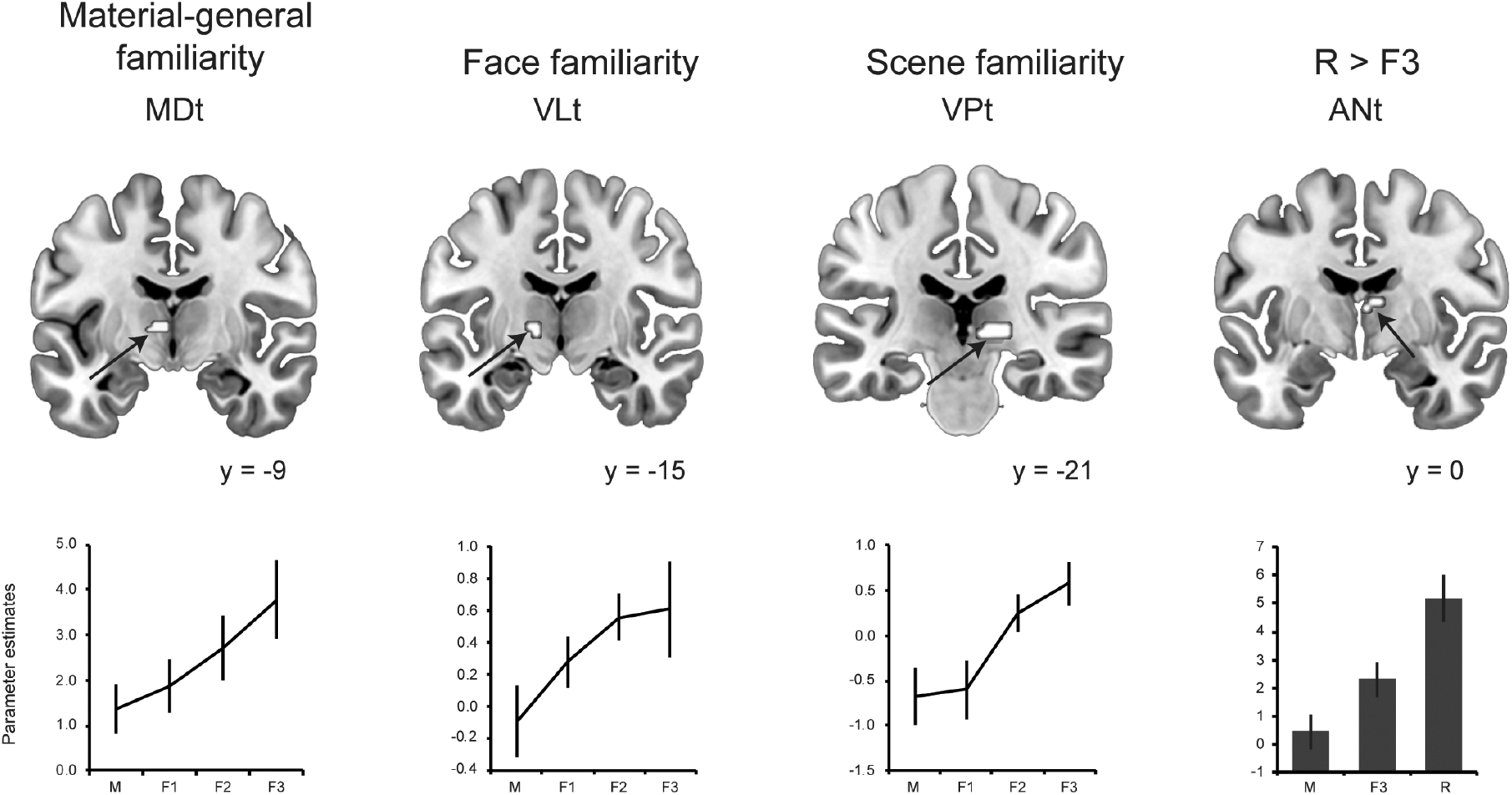
Familiarity and recollection effects in the thalamus. The MDt (mediodorsal thalamus) responded in a material-general way to familiarity across faces, objects and scenes (conjunction analysis), while the ventral lateral (VLt) and the ventral posterior (VPt) thalamic nuclei showed a material-specific familiarity response to faces and scenes (respectively). The anterior thalamus (ANt) showed greater response to recollection (R) than strong, accuracy matched, familiarity (F3). Activations are displayed at a voxel-wise *p* < 0.001 and are significant at a cluster-corrected family wise error (FWE) *p* < 0.05 determined via nonparametric permutations (all *t*s > 3.68). M = misses. Error bars show the standard error of the mean.

### The Familiarity Network: Material-specific and material-general effects

In order to identify the brain regions that make up the material-specific and material-general familiarity networks, familiarity responses for each stimulus type were modeled parametrically, exploring increases or decreases in activity across reported familiarity strength (see Methods). Material-general activity patterns were explored using a conjunction analysis of the parametric familiarity responses across the three types of stimulus. To explore material-specific activity patterns, each one of the parametric models for scenes, faces and objects was exclusively masked by the parametric models of the other two types of stimulus (see Methods). These analyses revealed the brain regions that respond to familiarity irrespective of stimulus type (conjunction), as well as those regions that uniquely respond to each of the three types of stimuli (parametric effects with exclusive masks). We especially target effects in the different thalamic regions, but whole brain effects are shown in Figure 1 and are reported in the Suppl. Results (see also Suppl. Tables 2 – 5).

### Thalamic familiarity and recollection effects

The thalamic regions revealed a combination of material-specific and material-general responses to familiarity as shown in Figure 2. A cluster within the MDt (extended into the ventrolateral nucleus; peak MNI: −6 −9 6) was found to respond to familiarity strength in a material-general way, while the right ventral posteromedial thalamus (VPt; including the pulvinar; peak MNI: 9 −21 −3) revealed activation selective to scene familiarity, and the left ventrolateral thalamus (VLt; peak MNI: −12 −15 0; 75 voxels) revealed a selective response to face familiarity. Activation produced by recollection (R) was contrasted with that produced by F3 (strong familiarity, matched in accuracy to R). Within the thalamus, a selective response to R versus F3 was found bilaterally in the ANt (peak MNI: −3 0 9 and 6 −8 12; Figure 2). The whole-brain activation data from this contrast are reported in Suppl. Table 6.

### Connectivity between thalamic regions and the MTL

A whole-brain psychophysiological interaction (PPI) analysis was used to explore the functional connectivity patterns between the active thalamic regions (reported above) and the structures of the MTL or the whole brain, as a function of reported familiarity. A significant PPI in the present study indicates that activity in a target region is modulated by, or co-varies with, activity in the seed area (thalamic regions) when participants engage in familiarity decisions for the three types of stimuli. For each seed region both positive and negative PPI connectivity was considered (see Methods), but significant negative PPI was only found with the VLt seed for face familiarity. The remaining PPIs, reported below, denote positive connectivity between the thalamic seed areas and the target regions. The identified target regions for the different PPI analyses (with the different seed regions) are reported in Suppl. Table 7 and Figure 3.

**Figure 3.**
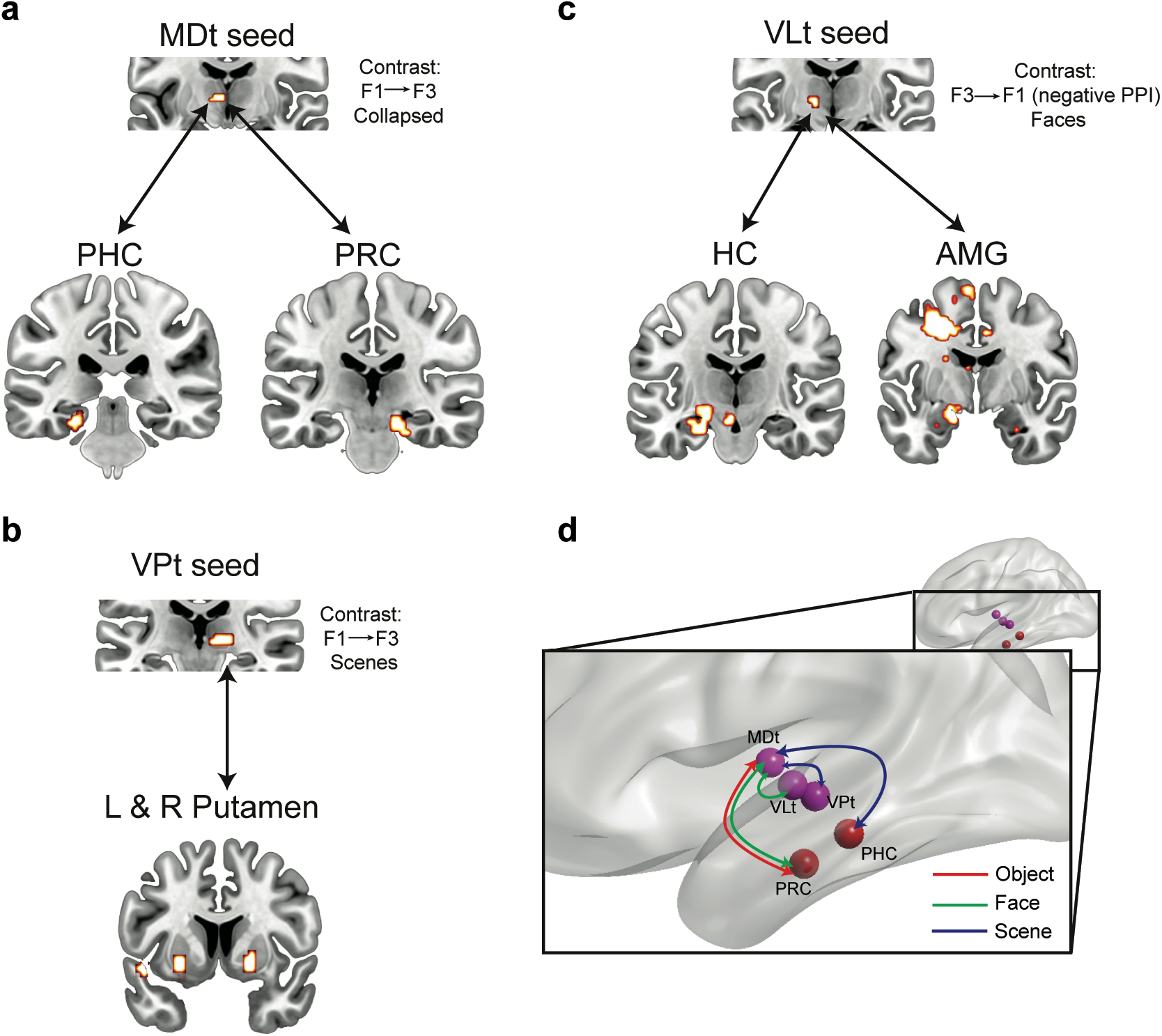
Functional connectivity analysis (PPI) using three thalamic seeds driven by increased familiarity strength (positive PPI; a and b) or decreased familiarity strength (negative PPI; c). Clusters of significant activation are displayed at a voxel-wise *p* < 0.001 and are significant at cluster-wise FWE *p* < 0.05 determined with nonparametric permutation testing (with 5000 permutations; all *t*s > 3.75). d) Schematic summary of intra-thalamic and thalamic-MTL functional connectivity patterns supporting familiarity memory. Material-specificity in the connectivity profiles is informed by the univariate and PPI analyses and correlational analysis reported in Suppl. Results and Suppl. Figure 1. MDt-PRC connectivity supports familiarity for objects and faces, while MDt-PHC connectivity supports familiarity for scenes. Within the thalamus, material-specific regions (VLt for faces and VPt for scenes) directly communicated with the MDt. Abbreviations: PHC = parahippocampal cortex; PRC = perirhinal cortex; HC = hippocampus; AMG = amygdala; MDt = mediodorsal thalamus; VLt = ventral lateral thalamus; VPt = ventral posterior thalamus (including pulvinar).

The PPI connectivity analysis using MDt (MNI: −6 −9 6) as the seed region for parametric increases in reported familiarity across the three types of stimuli, resulted in significant activations within the left PHC (BA 35; peak MNI: −24 −30 −12) and an area in the right posterior PRC (bordering PHC; BA 35; peak MNI: 21 −24 −21) extending to the midbrain/substantia nigra (MNI: 12 −21 −18; Figure 3a). Intra-thalamic connectivity with MDt was found with the left ventral lateral thalamus and the pulvinar (Suppl. Table 7 and Figure 3d). As reported above, in the whole-brain analysis, these thalamic areas were selectively sensitive to familiarity for faces and scenes, respectively and these findings may therefore signify the communication of material-specific familiarity information with the MDt, which was found to have a material-general role in familiarity memory.

The connectivity analysis using the VPt as the seed region (MNI: 9 −21 −3) for scene familiarity, identified regions within the bilateral dorsal striatum (putamen; Figure 3b) and the right insula (BA 13). No connectivity was identified with any MTL structures in this analysis. Finally, the connectivity analysis with LVt as the seed, did not show any significant positive PPI connectivity, but negative connectivity patterns were found. These were brain regions that increased their connectivity with the seed area when *decreased* levels of familiarity for faces were reported. These included the parahippocampal cortex (BA 35), the hippocampus and the amygdala within the MTL (Figure 3c), as well as the bilateral superior medial frontal gyrus (BA 6/10), the right precuneus (BA 7), the bilateral lingual gyrus and the left fusiform gyrus (BA 19).

The connectivity analysis using the ANt as the seed region (MNI: 6 −8 12) for R versus F3 did not produce any significant activations either in the MTL or in the whole brain.

## Discussion

In the current study, we explored the contribution of the different thalamic nuclei to memory and asked whether functional connectivity between the thalamic nuclei and the MTL is critical for recognition memory. For the first time, material-specific (ventral lateral and posterior nuclei) and material-general (MDt) thalamic regions were identified as supporting familiarity memory, while the anterior thalamus was found to support recollection when compared to equally strong familiarity. Finally, we also provide the first direct evidence that familiarity memory relies on a functional coupling between the thalamus (MDt) and the MTL (PRC and PHC) and the degree of this communication relates to the strength of reported familiarity. This provides strong evidence that this coupling directly supports familiarity memory. These findings are critical for our understanding of the key role played by the thalamus in recognition memory (Aggleton and Brown 1999; Aggleton et al. 2011; Carlesimo et al. 2015) and the importance of the thalamic-MTL interplay.

### The role of the thalamic nuclei in recognition memory and their dynamic interactions with the MTL

Material-specific familiarity-driven activity in the thalamus was found in the present study for face and scene stimuli. Face familiarity resulted in selective activation within the left VLt, whereas scene familiarity selectively activated a more posterior area of the thalamus including the ventral posteromedial and pulvinar nuclei (VPt). However, the MDt – extending laterally into the ventral (anterior and lateral) thalamus – responded to familiarity strength across all three stimulus types.

The MDt has previously been reported as belonging to a network of brain regions that supports familiarity memory (Aggleton and Brown 1999; Aggleton et al. 2011). Previous fMRI studies have shown that the MDt responds to increases in familiarity strength for object and scene stimuli (Montaldi et al. 2006; Kafkas and Montaldi 2014) and responds selectively to strong familiarity but not to equally strong recollection (Kafkas and Montaldi 2012; but see Pergola et al. 2013 for a different pattern at encoding). The current study additionally suggests that the MDt, unlike the MTL cortices, has a material-general role in familiarity-based recognition, as the hippocampus does for recollection-based recognition (Kafkas et al. 2017). Consistent with this suggestion, the MDt holds reciprocal anatomical connections with the anterior parahippocampal gyrus and the amygdala (Krettek and Price 1974; Aggleton and Mishkin 1984; Russchen et al. 1987; Goulet et al. 1998; Saunders et al. 2005), regions that were also found in our previous study (Kafkas et al. 2017) to be sensitive to stimulus familiarity for different types of stimulus. Therefore, it is possible that familiarity signals computed in the different MTL regions, for different stimulus types (see Kafkas et al., 2017), converge in MDt. Indeed, our connectivity analysis reinforces this suggestion as the MDt was found to be functionally connected with MTL structures, also responsible for supporting familiarity (PRC and PHC), and the degree of this connectivity determines the degree of familiarity confidence. Correlational analyses (Suppl. Results and Suppl. Figure 1) showed that a degree of material selectivity of the MTL-MDt functional coupling may be in operation, with the PRC-MDt coupling correlating with familiarity performance for objects and faces and the PHC-MDt coupling correlating with familiarity performance for scenes. Nevertheless, some caution is warranted interpreting this analysis because of the raised risk of false positives when small numbers of participants are used.

With the same paradigm (Kafkas et al., 2017), we have previously found that the amygdala (along with ERC) selectively responds to face familiarity (relative to familiar objects and scenes), whereas here we are reporting that connectivity between the MDt and the PRC is more critical for face familiarity. In contrast, no significant functional connectivity pattern was found between the amygdala and the MDt. However, the VLt that also selectively responded to face familiarity, was connected to the amygdala showing a negative PPI effect (Figure 3c), which indicates stronger connectivity when weaker feelings of familiarity occur (see also below). Therefore, the previous (from Kafkas et al., 2017) and current findings indicate that familiarity processing within the amygdala appears to be more local and certainly does not depend on the connectivity with thalamic regions. Instead functional connectivity with the MDt during familiarity decisions only occurs via the PRC, unless weak familiarity is detected which results in increased VLt-amygdala connectivity.

Intra-thalamic connectivity was also found between MDt and the face specific and scene specific thalamic regions similar to those identified in the main analysis (VLt and VPt). These findings suggest that different thalamic nuclei compute, or carry, familiarity signals depending on the type of stimulus that is processed, but these material-specific familiarity signals are selectively communicated with the MDt which has a material-general role in familiarity memory for pictorial stimuli (Figure 3d). This is further reinforced by the finding that only the MDt was functionally connected to the MTL cortices, while the other thalamic seeds (VLt and VPt) did not show increased connectivity with the MTL. Collectively, these findings further reinforce the hypothesis that the MDt is a structure of convergence of familiarity signals and potentially a critical hub of information integration supporting familiarity memory (Kafkas and Montaldi 2018).

Human lesion studies have provided mixed results with respect to the selectiveness of the MDt in supporting familiarity memory (Zoppelt et al. 2003; Cipolotti et al. 2008; Soei et al. 2008; Pergola et al. 2012), probably owing to the fact that most patients have damage that spreads into other thalamic nuclei or into parts of the extended recollection network, which may obscure selective familiarity deficits. A recent study, however, reported a face familiarity deficit with spared recollection in a patient with right MDt damage (Edelstyn et al. 2016). Yet another recent neuropsychological study reported both recollection and familiarity deficits for word stimuli in a group of MDt patients (Danet et al. 2017). Although the findings from these two neuropsychological studies appear incompatible, there are important methodological differences between them that should be considered when examining the specific role of the MDt. Most prominently, the type of stimulus used in these studies varied (faces were used in Edelstyn et al. and words in Danet et al.), which leaves open the possibility that the MDt contributes differently to pictorial and verbal familiarity memory.

When examining the role of the MDt in recognition memory as well as the extent to which this structure selectively contributes to familiarity, another important factor is its division into magnocellular (MDtm) and parvocellular (MDtp) portions (Russchen et al. 1987; Aggleton and Brown 1999). These regions have different connectivity patterns (e.g., Mitchell and Chakraborty 2013) and have been proposed to have different functional properties with only the MDtm portion connected to the MTL and subserving familiarity memory (Aggleton and Brown 1999; Zoppelt et al. 2003). In contrast, the MDtp, which does not connect to the MTL but mainly to the prefrontal cortex, has been found to be more critical for recall (Pergola et al. 2012). Although, the activations produced within the MDt in the present study appear to predominantly fall within the MDtm, its spatial resolution does not allow confident differentiation of whether the BOLD signal originates from either the MDtm or the MDtp. That said, the functional connectivity patterns characterizing the MDt in the present study, as an area that not only responds to familiarity but also connects to the MTL neocortices when familiarity decisions are involved, are more consistent with the proposal that the observed activation is predominantly located within the MDtm.

The anterior thalamus (ANt) was found to respond to recollection across all three kinds of stimulus, when this was contrasted with equally strong familiarity (F3). This finding is consistent with previous fMRI (Kafkas and Montaldi 2012; Pergola et al. 2013), neuropsychological (Clarke et al. 1994; Carlesimo et al. 2007; Pergola et al. 2012) and animal studies (Wilton et al. 2001) regarding the role of the ANt in selectively supporting recollection or other forms of associative memory. Similarly, in previous fMRI studies (Yonelinas et al. 2005; Cohn et al. 2009; Kafkas and Montaldi 2012), the hippocampus has also been shown to respond predominantly to recollection relative to strength-matched familiarity and, in recent work, this role was found to be material-general (Kafkas et al. 2017). Therefore, as with the hippocampus, the ANt seems to support recollection, regardless of the kind of stimulus that triggers it. The functional similarity between these structures can be attributed to the extensive anatomical connections characterizing the links between the ANt and the hippocampus (Aggleton et al. 2010). However, in the present study we failed to find increased functional connectivity between the two when recollection responses were modeled in the PPI analysis. This null finding is difficult to interpret, but it may relate to the smaller number of recollections reported in our paradigm, which encourages more familiarity-based recognition. Therefore, spontaneous recollection following a shallow encoding task (as used here), while sufficient to engage the hippocampus (see Kafkas et al. 2017) and the ANt more than strong familiarity, might require more experimental power (e.g., increased instances of recollection) to detect the functional connectivity between these two structures.

We turn now to ask what the specific role might be of the two thalamic regions that selectively respond to face and scene familiarity (VLt and VPt, respectively). The effects observed in VLt and VPt are novel and therefore future studies will be needed to explore and replicate whether these structures play a critical role in material-specific familiarity. That said, the selective familiarity effect in the ventrolateral thalamus for face familiarity, found in the present study, is highly consistent with prior observations. The potential contribution of the VLt to recognition memory has been reported in previous human lesion studies (Von Cramon et al. 1985; Zoppelt et al. 2003; Gold and Squire 2006; Soei et al. 2008). Furthermore, in an older study (Vilkki and Laitinen 1976), impaired face-matching performance was reported in a group of 38 patients following ventrolateral thalamotomy, while two more patients with ventral thalamic lesions were identified as having face-specific recognition impairments after longer retention intervals (48 hours) (Von Cramon et al. 1985).

The posterior thalamic nuclei and the pulvinar (VPt), which were found in the current study to have a selective role in scene familiarity, are considered to be a key part of the ‘visual thalamus’ and are densely interconnected with visual cortex, and with frontal, parietal and temporal regions (Shipp 2003; Saalmann and Kastner 2009; Arcaro et al. 2015). The pulvinar, in particular, constitutes a relay of cortico-cortical communication (Shipp 2003) and has been linked to tasks requiring visuospatial attention (Arend et al. 2008), goal-directed selection and visual binding (Snow et al. 2009; Wilke et al. 2013). Critically, this structure may play an important part in flexibly integrating visual information (Saalmann and Kastner 2009). Therefore, the role of the pulvinar in the integration and maintenance of relevant visual information is likely to contribute to memory performance (Vilkki 1978; Rotshtein et al. 2011) and our findings suggest that this role may be particularly critical when scene information is processed and retrieved. Although, both VLt and VPt were found to generate material-specific familiarity signals, neither of them was functionally connected to the MTL. We hypothesize that any possible communication between these thalamic structures and the MTL may be indirectly mediated through the MDt, whose functional connectivity with both the MTL and the VLt and VPt was found to underpin familiarity memory. However, due to the lack of direct evidence of mediation of connectivity from VLt/VPt to the MTL exclusively via MDt, further research is needed to directly test this hypothesis.

One notable exception to this positive connectivity profile is the *negative* connectivity characterizing the VLt and the MTL, including the hippocampus, the amygdala, and the PHC. The negative PPI found here, suggests that the VLt was functionally connected to the MTL structures more when weaker rather than stronger feelings of familiarity were reported. To further explore the reason for this effect, a further PPI analysis with the VLt seed was run for new faces (CR) versus weakly familiar faces (F1), which also indicated increased connectivity with the same MTL regions (Suppl. Table 8). Therefore, the functional connectivity between the VLt and the MTL prioritizes the more novel faces, perhaps reflecting the triggering of further encoding of face information, which may increase subsequent familiarity and/or recollection.

## Conclusions

In summary, these findings illustrate how different thalamic nuclei provide contrasting specialized support for memory; the MDt and the ANt are specialized to support familiarity and recollection, respectively, in a material-general way, whereas the ventral and posterior thalamic regions selectively support familiarity for particular kinds of visually complex stimuli; faces and scenes. Moreover, the MDt integrates familiarity signals from other thalamic regions in order to directly communicate this with the MTL cortices and the extent to which this communication occurs has a modulatory effect on the level of familiarity experienced.

## Acknowledgments

This work was supported by the Wellcome Trust (grant number: 094597/Z/10/Z). The authors declare no competing interests.

