## Supplementary material for "Thalamic-medial temporal lobe connectivity underpins familiarity memory": Suppl.

Alex Kafkas, Andrew R. Mayes & Daniela Montaldi
School of Biological Sciences, Division of Neuroscience & Experimental Psychology, University of Manchester, UK

#### Supplementary Results and Discussion

**Whole-brain familiarity effects**

Apart from the thalamic familiarity effects as described in the main text of the paper, material-specific and material-general effects were also found in other brain regions (for the effects within the MTL see Kafkas et al. (2017). The familiarity network comprised regions consistently responding to familiarity strength in a material-general way and included frontal, parietal and subcortical structures (Figure 1c and Suppl. Table 1). Middle and medial prefrontal regions corresponding to the dorsolateral and medial prefrontal cortex (BA 8/9), the anterior cingulate (BA 32) and the inferior frontal gyrus (BA 45/44) were found to increase their activity across familiarity strength consistently for all stimulus types. In the parietal lobe, the inferior parietal lobe including the angular gyrus (BA 40), the posterior cingulate cortex (BA 23/31) and the precuneus (BA 7) also responded to familiarity strength.

Areas showing selective and unique activations to scene familiarity (see Suppl. Table 2) included an area in the left middle frontal and orbitofrontal cortex (BA 10/11) and the right dorsal striatum (bilaterally) extended into the midbrain. Unique parametric responses to object familiarity (see Suppl. Table 3), exclusively masked by scene and face familiarity, were found in an area within the left dorsolateral prefrontal cortex (BA 6/8) and the left ventrolateral prefrontal cortex (BA 44/45/46). Importantly, parametric deactivations tracking familiarity strength within the fusiform gyrus also uniquely characterized object familiarity. Finally, a cluster in the right superior parietal lobe (BA 7), selectively tracked face familiarity (see Suppl. Table 4).

Most of the PFC regions (dorsolateral and medial prefrontal cortex (BA 8/9), the anterior cingulate (BA 32) and the inferior frontal gyrus (BA 45/44) were found to play a material-general role in familiarity-based recognition, encompassing all stimulus types (i.e., objects, faces and scenes; Figure 1c). However, two areas in the left dorsolateral PFC and the left ventrolateral PFC activated selectively to familiarity for object stimuli, whereas a frontopolar and orbitofrontal cortex (BA 10/11) area selectively responded to scene familiarity. These findings are consistent with previous fMRI studies (Wheeler and Buckner 2004; Yonelinas et al. 2005; Montaldi et al. 2006; Kafkas and Montaldi 2012, 2014) and with lesion studies reporting familiarity deficits after lateral (Duarte et al. 2005; Aly et al. 2011) or anterior/medial PFC damage (MacPherson et al. 2008). The inferior parietal lobe (including the angular gyrus), the precuneus and the posterior cingulate cortex were also found to respond to familiarity-based recognition across the three stimulus types. Only the right superior parietal lobe (BA 7) emerged as an area with a material-specific role; this was for familiarity for faces. It has been argued that the dorsal parietal lobe (which includes the superior parietal area) plays a role in familiarity-based recognition (Cabeza et al. 2008; Vilberg and Rugg 2008; Kim 2010). However, in the present study, consistent with previous fMRI evidence (Yonelinas et al. 2005; Montaldi et al. 2006; Kafkas and Montaldi 2012), the inferior parietal lobe was also found to respond to the degree of familiarity experienced and importantly, in a material-general manner. These PFC and parietal regions were not, however, selectively sensitive to familiarity as they were also activated when recollection was compared with strong familiarity (see Suppl. Table 5). This suggests that PFC and parietal areas play a role in processes that contribute to both familiarity and recollection, which is consistent with the argument that familiarity and recollection are also supported by some common neural and behavioural mechanisms.

**Does the connectivity between thalamus and MTL correlate with familiarity performance?**

We also explored whether any functional coupling between the thalamic regions and the MTL, as reported in the paper, contribute to familiarity performance. To explore this, the relationship between the degree of connectivity within the isolated regions (expressed as the PPI beta estimates) and behavioural familiarity performance was examined using (non-parametric) Spearman's rho correlations. We selected to use Spearman's correlation coefficients to overcome possible assumption violations from the use of parametric tests, such as Pearson's correlation coefficients when smaller samples are used. For the correlational analyses, familiarity performance was calculated using d' (the same analyses using Hits – FAs produced very similar results).

These analyses showed that the degree of MDt-PRC connectivity reliably correlated with familiarity performance (d') collapsed across the three types of stimuli (Suppl. Figure 1a; $r_s$ = 0.67, $p$ = 0.003). This was also true when the relationship was examined separately for object familiarity performance ($r_s$ = 0.64, $p$ = 0.005) and face familiarity performance ($r_s$ = 0.51, $p$ = 0.038) but not for scene familiarity performance ($r_s$ = 0.18, $p$ = 0.48; Suppl. Figure 1a). Overall, these effects indicate that the greater the degree of MDt-PRC connectivity the better the familiarity performance especially for objects and faces. Finally, the degree of connectivity between the MDt and the PHC did not reliably correlate with overall familiarity performance ($r_s$ = -0.07, $p$ = 0.79) or familiarity performance for objects ($r_s$ = -0.064, $p$ = 0.81) or faces ($r_s$ = -0.12, $p$ = 0.63), however, it significantly correlated with familiarity performance for scenes ($r_s$ =

0.53, $p$ = 0.028; Suppl. Figure 1b). A comparison between the correlations for MDt-PHC connectivity with performance for the different types of stimuli (using Fisher's z transformation for overlapping correlations in dependent groups, see Myers and Sirois 2006; Diedenhofen and Musch 2015) showed that these differences in correlations between scenes and objects ($Z$ = 2.38, $p$ = 0.017) and scenes and faces ($Z$ = 2.32, $p$ = 0.02) were statistically significant. This confirms that the correlation between MDt-PHC connectivity and familiarity performance was stronger and selectively significant for scenes (but not objects and faces). Overall, these correlation analyses indicate a degree of material-specificity characterizing the connectivity between MDt and the PRC/PHC in familiarity. MDt-PRC connectivity correlated with object and face familiarity, while MDt-PHC connectivity correlated with scene familiarity.

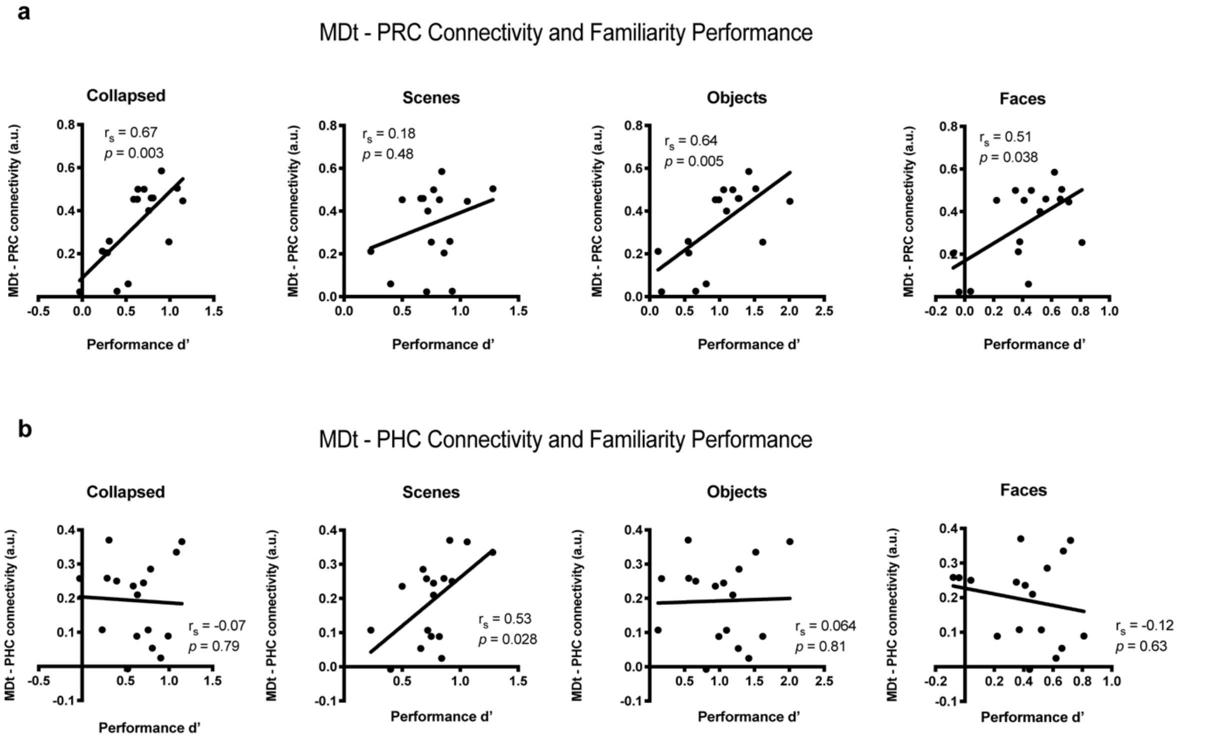


**Supplementary Figure 1.** Relationship between the degree of connectivity and familiarity performance. Degree of connectivity between MDt and PRC (a) or PHC (b) and familiarity performance collapsed and separately for the three types of stimuli. MDt-PRC connectivity reliably correlated with familiarity performance for objects and faces. MDt-PHC connectivity reliably correlated with familiarity performance for scenes only. Spearman's rho correlation coefficients ($r_s$) are reported.

### Supplementary Tables

**Supplementary Table 1.** Proportion of trials (standard deviations in the parentheses) across three types of stimuli (scenes, objects and faces) for the different response outcomes in the recognition task

| | Hit F1 | Hit F2 | Hit F3 | Hit R | CR | M | FA F1 | FA F2 | FA F3 | FA R |
|---|---|---|---|---|---|---|---|---|---|---|
| **Scenes** | 0.24 (0.14) | 0.18 (0.08) | 0.24 (0.17) | 0.11 (0.06) | 0.69 (0.10) | 0.23 (0.10) | 0.17 (0.08) | 0.07 (0.04) | 0.04 (0.03) | 0.03 (0.01) |
| **Objects** | 0.14 (0.09) | 0.22 (0.10) | 0.37 (0.19) | 0.15 (0.16) | 0.71 (0.23) | 0.12 (0.12) | 0.18 (0.16) | 0.07 (0.06) | 0.03 (0.03) | 0.01 (0.01) |
| **Faces** | 0.37 (0.15) | 0.23 (0.13) | 0.12 (0.11) | 0.02 (0.02) | 0.46 (0.26) | 0.26 (0.20) | 0.32 (0.17) | 0.16 (0.13) | 0.06 (0.08) | |
| **Collapsed** | 0.25 (0.11) | 0.21 (0.07) | 0.23 (0.14) | 0.10 (0.07) | 0.60 (0.21) | 0.21 (0.12) | 0.26 (0.17) | 0.10 (0.07) | 0.03 (0.04) | 0.01 (0.01) |

*Note:* F1= weak familiarity, F2 = moderate familiarity, F3 = strong familiarity, R = recollection, CR = correct rejections, M = misses, FA = false alarms.

**Supplementary Table 2.** Monotonic increases in activity across familiarity strength shared across the three types of stimuli

| Side | Region | No. of Voxels | ~BA | MNI x y z | *T*-value |
|---|---|---|---|---|---|
| L | Superior frontal gyrus | 311 | BA 8/6 | -21 24 57 | 5.33 |
| L | Superior medial frontal gyrus | | BA 8/9/32 | -9 30 54 | 4.99 |
| L | Middle frontal gyrus and orbitofrontal gyrus | 236 | BA 10/46/47 | -30 60 3 | 5.22 |
| L | Inferior parietal lobe | 170 | BA 40 | -42 -54 54 | 5.11 |
| L | Precuneus | 161 | BA 7 | -6 -60 39 | 4.93 |
| L | Posterior cingulate/ retroplenial | | BA 23/31 | 0 -39 24 | 4.10 |
| L | Inferior frontal gyrus | 145 | BA 45/46 | -45 27 33 | 4.41 |
| L | Caudate | 19 | | -12 15 0 | 4.16 |
| R | Superior parietal lobe | 32 | BA 7 | 24 -72 57 | 4.15 |
| L | Thalamus (MDt) | 18 | | -6 -9 6 | 3.87 |

*Note:* Activations are cluster FWE-corrected for multiple comparisons at $p < 0.05$ determined via nonparametric permutations ($t$s > 3.68)

**Supplementary Table 3.** Material-general and material-specific monotonic increases in activity across familiarity strength for scenes

| Side | Region | No. of Voxels | ~BA | MNI x y z | T-value |
|---|---|---|---|---|---|
| ***Material-general effects*** | | | | | |
| L | Superior medial frontal gyrus | 315 | BA 6/32 | -6 27 57 | 5.52 |
| L | Superior frontal gyrus | | BA 6/8 | -33 18 57 | 5.49 |
| L | Inferior parietal lobe | 102 | BA 40 | -36 -63 51 | 4.58 |
| L | Precuneus | 67 | BA 7 | -3 -57 39 | 4.55 |
| L | Posterior cingulate cortex | | BA 31 | 0 -42 30 | 3.87 |
| L | Angular gyrus | 38 | BA 39 | -48 -60 24 | 4.10 |
| L | Caudate | 14 | | -6 15 6 | 4.04 |
| L | Middle frontal gyrus | 41 | BA 9 | -45 30 33 | 3.83 |
| ***Material-specific effects*** | | | | | |
| R | Perirhinal/entorhinal cortex | 9 | BA 28/35 | 15 -6 -27 | 4.15 |
| R | Putamen/Lentiform nucleus | 7 | | 30 -3 -3 | 3.67 |
| L | Middle Frontal Gyrus | 392 | BA 10 | -27 60 6 | 5.41 |
| L | Orbitofrontal cortex | | BA 10/11/47 | -30 45 -3 | 4.87 |
| R | Thalamus, midbrain and lentiform nucleus | 206 | | 9 -21 -3 | 4.39 |
| R | Parahippocampal Cortex | | BA 35 | 24 -30 -12 | 3.80 |

*Note:* Activations are cluster FWE-corrected for multiple comparisons at $p < 0.05$ determined via nonparametric permutations ($t$s > 3.68)

**Supplementary Table 4.** Material-general and material-specific monotonic increases and decreases in activity across familiarity strength for objects

| Side | Region | No. of Voxels | ~BA | MNI x y z | T-value |
|---|---|---|---|---|---|
| ***Material-general effects*** | | | | | |
| L | Inferior parietal lobe | 155 | BA 40/39 | -42 -60 51 | 4.14 |
| L | Angular gyrus | | BA 40 | -48 -63 39 | 3.71 |
| L | Orbitofrontal cortex | 37 | BA 47 | -48 36 0 | 3.92 |
| ***Material-specific activations*** | | | | | |
| L | Superior frontal gyrus | 179 | BA 6/8 | -15 24 54 | 4.53 |
| L | Inferior frotnal gyrus | 200 | BA 44/45/46 | -54 18 21 | 4.00 |
| ***Material-specific deactivations*** | | | | | |
| R | Fusiform gyrus | 13 | BA 37 | 33 -66 -9 | 3.76 |
| R | Fusiform gyrus | 21 | BA 37 | 45 -54 -9 | 3.73 |
| L | Lingual gyrus | 6 | BA 18 | -18 -78 -6 | 3.73 |

*Note:* Activations are cluster FWE-corrected for multiple comparisons at $p < 0.05$ determined via nonparametric permutations ($t$s > 3.68)

**Supplementary Table 5.** Material-general and material-specific monotonic increases in activity across familiarity strength for faces

| Side | Region | No. of Voxels | ~BA | MNI x y z | T-value |
|---|---|---|---|---|---|
| ***Material-general effects*** | | | | | |
| L | Superior parietal lobe | 24 | BA 7 | -33 -63 48 | 3.81 |
| L | Middle frontal gyrus | 42 | BA 6/8 | -27 18 57 | 3.92 |
| L | Middle cingulate gyrus | 17 | BA 32 | -3 24 36 | 3.74 |
| ***Material-specific effects*** | | | | | |
| L | Lentiform nucleus | 75 | | -15 0 6 | 4.53 |
| L | Caudate | | | -12 12 0 | 3.82 |
| L | Thalamus | | | -12 -15 0 | 3.79 |
| L | Amygdala / Entorhinal cortex | 35 | BA 34 | -21 0 -24 | 4.05 |
| R | Superior Parietal lobe | 27 | BA 7 | 15 -69 57 | 4.00 |

*Note:* Activations are cluster FWE-corrected for multiple comparisons at $p < 0.05$ determined via nonparametric permutations ($t$s > 3.68)

**Supplementary Table 6.** Whole-brain activity for R versus F3 responses

| Side | Region | No. of Voxels | ~BA | MNI x y z | *T*-value |
|---|---|---|---|---|---|
| L&R | Precuneus and posterior cingulate | 396 | BA 7 | 0 -51 39 | 7.49 |
| L&R | Caudate nucleus | 94 | | -12 21 0 | 5.34 |
| L&R | Thalamus | 40 | | -3 0 9 | 5.06 |
| | | | | 6 -8 12 | |
| L | Anterior Cingulate | 257 | BA 32 | -9 45 9 | 6.16 |
| L | Superior medial frontal gyrus | 277 | BA 10 | -9 60 6 | 6.01 |
| L | Middle frontal gyrus | 141 | BA 10 | -21 18 45 | 5.84 |
| R | Inferior frontal gyrus | 195 | BA 47 | 51 24 -6 | 5.74 |
| R | Superior temporal gyrus | 63 | BA 22 | 54 12 -3 | 5.36 |
| L | Inferior frontal gyrus and Insula | 182 | BA 47 | -51 24 -3 | 5.39 |
| L | Superior temporal gyrus | 38 | BA 22 | -51 3 -9 | 4.07 |
| L | Middle temporal gyrus | 169 | BA 21 | -60 -54 3 | 5.29 |
| R | Middle temporal gyrus | 62 | BA 21 | 63 -45 6 | 5.23 |
| R | Superior medial frontal gyrus | 66 | BA 6 | 6 12 66 | 5.12 |
| L | Superior medial frontal gyrus | | | -6 12 63 | 4.21 |
| L | Hippocampus | 21 | | -15 -30 -6 | 4.43 |

*Note:* Activations are cluster FWE-corrected for multiple comparisons at $p < 0.05$ determined via nonparametric permutations (*t*s > 3.90)

**Supplementary Table 7.** Brain regions showing functional (PPI) connectivity with thalamic seeds as a function of reported familiarity strength

| Side | Region | Extent (voxels) | ~BA | MNI (x y z) | T-value |
|---|---|---|---|---|---|
| **Positive PPI with MDt (collapsed stimuli)** | | | | | |
| **L** | Inferior Parietal Lobe/Angular gyrus | 150 | BA39/40 | -39 -54 36 | 5.57 |
| **R** | Angular gyrus extending to Middle Occipital gyrus | 65 | BA39 | 33 -66 33 | 4.56 |
| **R** | Globus pallidus | 9 | | 12 0 -3 | 3.97 |
| **R** | Perirhinal cortex (extending to midbrain) | 37 | BA35 | 21 -24 -21 | 4.56 |
| **L** | Parahippocampal cortex | 8 | BA35 | -24 -30 -12 | 3.8 |
| **L** | Thalamus (pulvinar) | 11 | | -21 -30 12 | 3.8 |
| **L** | Ventral Lateral thalamus | 8 | | -9 -12 0 | 3.79 |
| **Positive PPI with VPt (scenes)** | | | | | |
| **L** | Insula & Putamen | 62 | BA 13/47 | -27 21 -9 | 5.05 |
| **R** | Putamen | 23 | | 18 6 -6 | 5.03 |
| **Negative PPI with VLt (faces)** | | | | | |
| **L** | Superior medial frontal gyrus | 182 | BA6 | -30 -6 42 | 6.43 |
| **R** | Superior medial frontal gyrus | 30 | BA10 | 12 54 6 | 5.40 |
| **L** | Parahippocampal cortex, hippocampus and amygdala | 156 | BA35/28 | -21 -24 -21 | 5.11 |
| **R** | Precuneus | 93 | BA7 | 15 -57 24 | 5.84 |
| **R** | Precuneus | 225 | BA7 | 6 -45 60 | 5.34 |
| **R** | Lingual gyrus | 159 | BA19 | 6 -87 -6 | 4.91 |
| **L** | Lingual gyrus and Fusiform gyrus | 75 | BA19 | -27 -84 -3 | 4.97 |

*Note:* Activations are cluster FWE-corrected for multiple comparisons at $p < 0.05$ determined via nonparametric permutations ($t$s > 3.75)

**Supplementary Table 8.** Brain regions showing functional (PPI) connectivity with VLt for new faces (CR) versus weakly familiar faces (F1)

| Side | Region | Extent (voxels) | ~BA | MNI (x y z) | T-value |
|---|---|---|---|---|---|
| **Positive PPI with VLt ($CR_{faces}$ > $F1_{faces}$)** | | | | | |
| **L** | Superior medial frontal gyrus | 187 | BA6 | -18 -6 42 | 7.40 |
| **L** | Parahippocampal cortex, hippocampus and amygdala | 166 | BA35/28 | -15 -24 -9 | 6.20 |
| **R** | Precuneus | 250 | BA7 | 15 -42 42 | 5.80 |
| **L** | Fusiform gyrus | 167 | BA19 | -15 -87 0 | 5.67 |

*Note:* Activations are cluster FWE-corrected for multiple comparisons at $p < 0.05$ determined via nonparametric permutations (*t*s > 3.75)

**Supplementary Table 9.** Brain regions showing functional (PPI) connectivity with thalamic seeds as a function of reported familiarity strength using 3mm thalamic seeds

| Side | Region | Extent (voxels) | ~BA | MNI (x y z) | T-value |
|---|---|---|---|---|---|
| **Positive PPI with MDt (collapsed stimuli)** | | | | | |
| **R** | Perirhinal cortex (extending to midbrain) | 72 | BA35 | 21 -24 -21 | 5.04 |
| **L** | Inferior Parietal Lobe/Angular gyrus | 91 | BA39/40 | -39 -54 36 | 5.01 |
| **L** | Orbitofrontal cortex | 36 | BA11 | -27 45 -9 | 4.80 |
| **L** | Ventral Lateral thalamus | 20 | | -9 -12 3 | 4.54 |
| **L** | Thalamus (pulvinar) | 34 | | -21 -30 12 | 4.02 |
| **L** | Putamen | 10 | | -24 6 6 | 3.78 |
| **Positive PPI with VPt (scenes)** | | | | | |
| **L** | Insula & Putamen | 75 | BA 13/47 | -27 23 -9 | 5.05 |
| **R** | Putamen | 20 | | 18 6 -6 | 4.92 |
| **Negative PPI with VLt (faces)** | | | | | |
| **L** | Parahippocampal cortex, hippocampus and amygdala | 165 | BA35/28 | -21 -24 -21 | 6.20 |
| **L** | Superior medial frontal gyrus | 152 | BA6 | -30 -5 42 | 6.11 |
| **R** | Superior medial frontal gyrus | 45 | BA10 | 12 54 6 | 5.50 |
| **R** | Precuneus | 120 | BA7 | 9 -66 45 | 5.45 |
| **R** | Precuneus | 150 | BA7 | 6 -45 60 | 5.20 |
| **R** | Lingual gyrus | 160 | BA19 | 6 -87 -6 | 5.01 |
| **L** | Fusiform gyrus | 65 | BA19 | -27 -69 -6 | 4.89 |

*Note:* Activations are cluster FWE-corrected for multiple comparisons at $p < 0.05$ determined via nonparametric permutations ($t$s > 3.75)

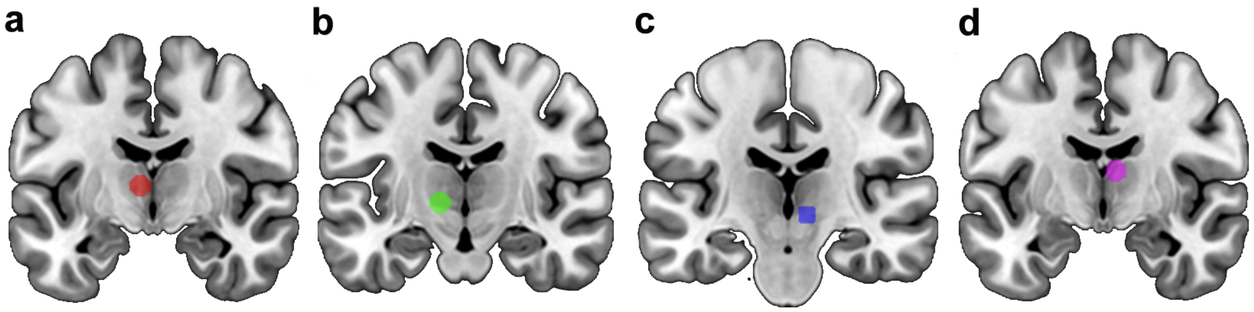


**Supplementary Figure 2.** Size and location of the thalamic seeds used in the psychophysiological interaction (PPI) analysis. The selection of the seeds was informed by the activations in the thalamus produced in the main parametric analysis for familiar stimuli (a,b and c) and the R > F3 contrast (d) as reported in the manuscript and shown in Figure 2. a) mediodorsal thalamus, MDt (red sphere, MNI: -6 -9 6); b) ventrolateral thalamus, VLt (green sphere, MNI: -12 -15 0); c) ventral posteromedial thalamus, VPt (blue sphere, MNI: 9 -21 -3); d) anterior thalamus, ANt (violet sphere, MNI: 6 -8 12).
